# Non-invasive fecal DNA yields whole genome and metagenomic data for species conservation

**DOI:** 10.1101/2022.08.16.504190

**Authors:** A. de Flamingh, Y. Ishida, P. Pečnerová, S. Vilchis, H.R. Siegismund, R.J. van Aarde, R.S. Malhi, A.L. Roca

## Abstract

Non-invasive biological samples benefit studies that investigate rare, elusive, endangered, and/or dangerous species. Integrating genomic techniques that use non-invasive biological samples with advances in computational approaches can benefit and inform wildlife conservation and management. Here we present a molecular pipeline that uses non-invasive fecal DNA samples to generate low- to medium-coverage genomes (e.g., >90% of the complete nuclear genome at 6X coverage) and metagenomic sequences, combining in a novel fashion widely available and accessible DNA collection cards with commonly used DNA extraction and library building approaches. DNA preservation cards are easy to transport and can be stored non-refrigerated, avoiding cumbersome and/or costly sample methods. The genomic library construction and shotgun sequencing approach did not require enrichment or targeted DNA amplification. The utility and potential of the data generated by this pipeline was demonstrated by the application of genome-scale analysis and metagenomics to zoo and free-ranging African savanna elephants (*Loxodonta africana*). Fecal samples collected from free-ranging individuals contained an average of 12.41% (5.54-21.65%) endogenous elephant DNA. Clustering of these elephants with others from the same geographic region was demonstrated by a principal component analysis of genetic variation using nuclear genome-wide SNPs. Metagenomic analyses generated compositional taxon classifications that included *Loxodonta*, green plants, fungi, arthropods, bacteria, viruses and archaea, showcasing the utility of our approach for addressing complementary questions based on host-associated DNA, e.g., pathogen and parasite identification. The molecular pipeline presented here extends applications beyond what has previously been shown for target-enriched datasets and contributes towards the expansion and application of genomic techniques to conservation science and practice.

## Introduction

Non-invasive biological samples benefit studies that investigate rare and elusive (Ferreira et al., 2018; Franklin et al., 2019), endangered (Ang et al., 2020) and/or dangerous species (Bellemain and Taberlet, 2004; Mondol et al., 2009). Non-invasively collected samples are obtained without directly interacting with animals, so that collection of samples do not impact the wellbeing of the animal from which it is collected (Taberlet et al., 1999; Sappington, 2019; Lefort et al., 2019). Non-invasive samples allow researchers to increase sample representation, by increasing geographic distribution and/or taxonomic diversity, or by substituting for physical sample collection procedures that may be arduous, expensive, or potentially harmful to the animal (e.g., chemical immobilization). The use of non-invasive samples has been complemented by advances in molecular techniques (Andrews et al., 2021) that progressively allow for smaller quantities of sample (and associated DNA template) to be sufficient for complex molecular analyses (Glocke and Meyer, 2017; Rohland et al., 2018; Xavier et al., 2021). Wildlife conservation genomics, the application of genomic techniques to inform conservation and management of species (Allendorf et al., 2010; Supple and Shapiro, 2018; Hohenlohe et al., 2021), can benefit from these technological advances, especially when used in synergy with non-invasive biological samples (Andrews et al., 2021; Hohenlohe et al., 2021). For example, combining non-invasive sampling with molecular analyses can benefit research and management of endangered species that may be difficult to sample due to low population numbers (Palomares et al., 2002; Baker et al., 2013; Carroll et al., 2018), or where invasive sampling can impact behavior and/or sociality (DelGiudice et al., 2001; Pelletier et al., 2004; Arnemo et al., 2006; Becciolini et al., 2019).

Here we present a molecular pipeline that combines established DNA collection, extraction and sequencing protocols in an innovative manner, allowing for the use of non-invasive fecal DNA samples to generate low-to medium-coverage whole genome and metagenomic sequences. In this study we aimed to combine the following: (1) a simplified and accessible DNA collection and extraction protocol that uses a standard DNA preservation card, which does not require refrigeration and is easy to transport and store; (2) a protocol for genomic library construction and shotgun sequencing that does not require enrichment or targeted DNA amplification; and (3) evaluate the utility and potential of the data generated by this approach through the application of genome-scale analysis and metagenomics of zoo and free-ranging elephants in their native environments.

Elephants are ideal candidates for testing a non-invasive approach because invasive collection of their samples can be unsafe and expensive(Jacobson et al., 1988; Kock et al., 1993), and they occur across very large and sometimes difficult to access geographic areas (Gray et al., 2014). Non-invasive samples have been used extensively to study elephants, e.g., to establish relatedness and demography (Munshi-South, 2011), investigate hybridization between forest and savanna elephants(Bonnald et al., 2021), study population structure and gene flow (de Flamingh et al., 2018), and estimate population size (Gray et al., 2014). We used samples from zoo individuals to investigate DNA content and preservation relative to dung freshness and verified the practicality and effectiveness of our pipeline in free-ranging elephants from five localities in South Africa.

## Methods and Materials

### Study design

We tested the use of DNA preservation cards (paper treated with chemicals that prevent microbial growth) in zoo elephants using a time-series of repeated samples (Figure 1). Three independent sets of samples were used (Supplementary Table 1); one set of 24 fecal samples from 6 elephants under human care from two zoos in the USA (Dataset 1), a set of 13 fecal DNA samples from 13 free-ranging elephants in five geographic localities in South Africa (Dataset 2), and a third set of reference samples from known geographic locations and for which genomic data had been previously generated from high quality tissue samples (Dataset 3).

**Figure 1:**
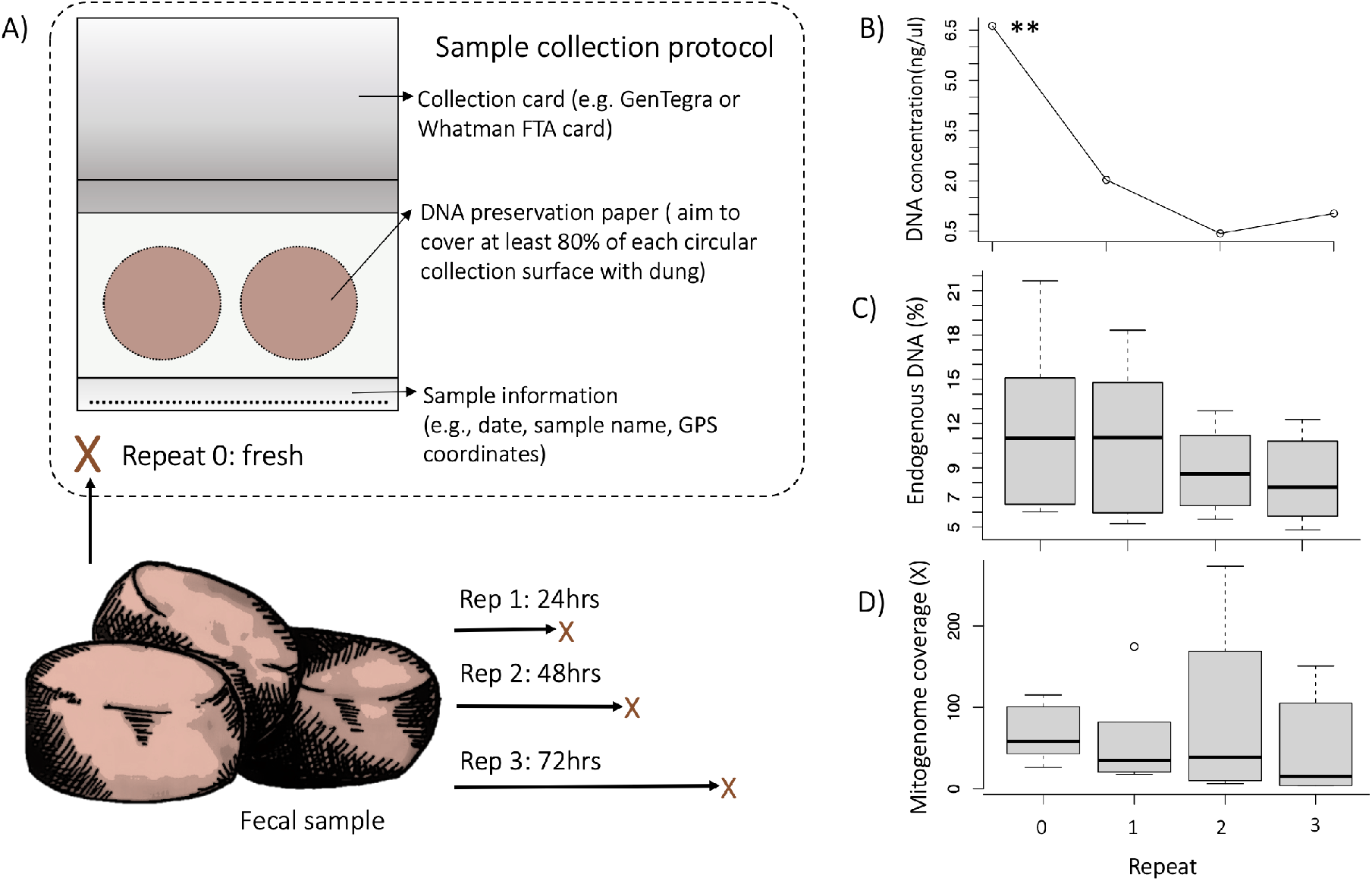
Experimental setup of replicate sampling. Assessment of the DNA concentration, endogenous DNA content (% reads mapping to the nuclear reference genome LoxAfr4.0) and mitogenome coverage (the average X-fold number of reads mapped at any location across the mitogenome) the comparison of repeated samples that were collected over time from the same dung bolus. A) Repeat samples were collected from dung immediately after defecation (Rep 0) and subsequently at 24hrs (Rep 1), 48hrs (Rep 2) and 72hrs (Rep 3) post defecation. Each sample was collected on an Ahlstrohm GenSaver 2.0™ card using both collection surfaces (indicated with circles). B) There was a significant decrease in DNA concentration as time after defecation increased (rep. measure ANOVA p = 0.0052). The percentage of endogenous elephant DNA also decreased as time after defecation increased (C), as did the mitogenome coverage (D).

For Dataset 1, samples were collected using the GeneSaver2™ Card (see Supplementary File 2 - DNA collection protocol) from three elephants housed at Jacksonville Zoo and Gardens, Florida, and three elephants from Dallas Zoological Gardens, Texas. For each elephant, four repeated samples were collected over time from the same dung bolus to quantify DNA degradation and to estimate how dung freshness may impact target-species DNA preservation. Repeated samples were collected from the same dung bolus (Figure 1) over 3 days: immediately after defecation (Rep 0), and then at 24hrs (Rep 1), 48hrs (Rep 2), 72hrs after defecation (Rep 3).

For Dataset 2, samples from 13 free-ranging elephants were collected from 5 geographic areas within South Africa (Supplementary Table 1; Supplementary Figure 1). Collectors targeted fresh dung, or dung that still had wet mucus (see Supplementary File 2 - DNA collection protocol).

Dataset 3 included 24 high-quality DNA samples that were geographically provenanced. These samples were used as references to evaluate the data that we generated in this study. This geo-referenced dataset included genomic data from four samples from Kruger National Park, South Africa (Pečnerová et al., *in prep*), one sample from Kenya and one sample from Kruger National Park (Palkopoulou et al., 2018), and 18 samples from Gorongosa National Park, Mozambique (Campbell-Staton et al., 2021; Supplementary Table 1; Supplementary Figure 1).

### Fecal DNA collection, extraction, and genomic library construction

The molecular pipeline used DNA preservation cards, which have also been used previously for conservation genomic (Régnier et al., 2011; Ward et al., 2019; Morrison et al., 2020) and microbiome research (Song et al., 2016; Yarlagadda et al., 2022). Preliminary testing compared two types of DNA preservation cards, the Whatman Flinders Technology Associates (FTA) card (Whatman plc, Maidstone, United Kingdom) and the Ahlstrom GenSaver 2.00™ card (GenTegra, Pleasanton, USA). Both types of cards resulted in similar extracted DNA quantity and quality as quantified using Qubit DNA quantitation and based on targeted PCR amplification of a 500bp elephant mitochondrial DNA fragment (Nyakaana and Arctander, 1999). We used only Ahlstrom GenSaver 2.00™ cards for subsequent DNA collection. Ahlstrom GenSaver 2.00™ cards are paper cards that have been treated with a chemical to minimize environmentally induced degradation and the growth of microorganisms, allowing for collection, transport, and long-term preservation of DNA from biological samples at ambient temperature (Ahlstrom Corporation, Helsinki, Finland). Fecal DNA was collected (Supplementary Figure 2) using a standardized collection protocol (Supplementary File 2 – DNA collection protocol). For free-ranging elephants, samples represented individuals from different herds, geographic locations, and different times of the day.

DNA was extracted using the QIAamp^®^ PowerFecal^®^ Pro DNA Kit (Qiagen, Ann Arbor, USA). Modifications to the standard DNA extraction protocol included the use of approximately one quarter of the collection card as the starting material (Supplementary Figure 2), after which samples were vortexed for 10 minutes. A maximum of 4 samples were processed per round of extraction. The final DNA elution step was repeated twice, each time using 50μl of Elution Buffer, resulting in a final DNA elute volume of 100μl.

Extracted DNA concentration was quantified using the Qubit Broad-Range dsDNA platform (ThermoFisher Scientific, Waltham, USA) and DNA concentration was compared across repeated samples (Dataset 1) using a repeated measures ANOVA in R (Core Team R, 2013) and a Bonferroni correction (source code available at github.com/adeflamingh). DNA concentration and endogenous content (see details below) were compared between fresh (repeat 0) samples from zoo individuals (Dataset 1) and samples from free-ranging individuals (Dataset 2) using a Welch Two Sample t-test. In addition, each repeat (1, 2, and 3) of Dataset 1 was also compared with Dataset 2 and a Benjamini-Hochberg correction was applied to account for multiple comparisons (Benjamini and Hochberg, 1995) in the program R (GitHub code).

Genomic libraries were constructed using a tagmentation library building approach as part of the Illumina DNA Prep Kit (ThermoFisher Scientific, Waltham, USA) and using the protocol described by (Yarlagadda et al., 2022). Repeats two and three of one of the zoo individuals did not show evidence of successful DNA extraction based on Qubit quantitation and amplification of the shorter 500bp mitochondrial sequence. We therefore did not build genomic libraries for those two repeat samples. To account for possible contamination, we also included negative control libraries with all rounds of sample processing. These libraries were constructed using blank GenSaver 2.00™ cards and the same reagents that were used for the fecal DNA libraries. We used IDT for Illumina DNA/RNA Unique Dual Indexes (Catalog number 20027213, Illumina, San Diego, Unites States), and shotgun-sequenced the pooled samples at the Roy. J. Carver Biotechnology Center, University of Illinois at Urbana-Champaign (UIUC), USA. Genomic library fragment size distribution was assessed by means of AATI (Advanced Analytical Technologies, Inc. Fragment Analyzer). Samples were sequenced in three independent runs (Supplementary Table 2, Supplementary Table 3) using the Illumina NovaSeq 6000 sequencing platform; the first two rounds of sequencing included the zoo, free-ranging and control libraries, and the third round included repeated sequencing (re-sequencing) of libraries of three individuals to increase depth and breadth of genome coverage (see below, Extended genomic analysis and prospective applications for details).

### Bioinformatic analyses

#### Quality control and genome alignment

Bioinformatic analysis used the Biocluster2 high-performance computing system at the Carl R. Woese Institute for Genomic Biology, UIUC, USA. Samples were de-multiplexed and the reads trimmed to a minimum length of 25bp using the program FastP v.0.19.6 (Chen et al., 2018). Reads were aligned to a reference African savanna elephant mitochondrial genome (mitogenome; GenBank accession number JN673264, Brandt et al., 2012) and nuclear genome (LoxAfr4.0; Meyer et al., 2017) using bowtie2 (Langmead and Salzberg, 2012) and BWA (Li and Durbin, 2010). Both alignment programs resulted in a similar number of reads mapping to mitogenome and nuclear genome reference sequences, and we used alignment files that were generated using the BWA-mem algorithm and default parameters in BWA for subsequent analysis. Alignments were transformed into BAM format using SAMtools v. 1.1 (Li et al., 2009), and filtered to remove unmapped reads and reads with a minimum alignment quality score less than 30. Filtered BAM files were then sorted and indexed, with PCR duplicates marked and removed with the Picard Toolkit v. 2.10.1 (Picard Toolkit, 2019, Broad Institute). Genome alignment statistics were calculated for each individual sample; we calculated the breadth (%) of each complete genome covered by reads; and the depth of coverage (X-fold) as the average number of reads mapping to each position on the complete genome using SAMtools (source code available at github.com/adeflamingh). The genomic percentage of endogenous elephant DNA was calculated as the fraction of reads mapping to the elephant nuclear genome (LoxAfr 4.0) to the total number of reads for a sample.

#### Metagenomic analysis and taxonomic classification

Unprocessed fastQ files were quality filtered and trimmed using Trimmomatic (Bolger et al., 2014): we removed adapters, leading and tailing low quality bases, and “N” bases with quality below 3; we scanned the read with a 4-base-wide sliding window, cutting reads when the average quality per base dropped below 15; to increase specificity for our metagenomic classification, we removed reads that were less than 50 bases long after trimming and only reads that were properly paired were retained for subsequent metagenomic analysis.

We used Kraken2 (Wood et al., 2019) to classify reads into taxonomic units. Kraken2 is a metagenomic sequence classifier that uses an ultrafast k-mer based approach to classify and assign taxonomic labels to short DNA reads (Wood et al., 2019). We compiled a custom search database to which reads were compared using the “kraken2-build” command. In addition to the African savanna elephant genome (LoxAfr 4.0,Meyer et al., 2017), our custom database included the RefSeq libraries archaea, bacteria, plasmid, viral, human, fungi, plant, and protozoa, the NCBI non-redundant nucleotide database, and UniVec_Core (NCBI database of vector, adapter, linker, and primer sequences). We concatenated pairs of reads together using the “--paired” function in Kraken2, parallelized the classification to run using 24 computing threads, and used a confidence scoring threshold of 0.05 to minimize erroneous classification and false positive rates (see Supplementary Figure 3 for details on confidence scoring threshold value). We also included the option for generating a summary report “—report” which was used estimate proportional read composition for comparison across repeated samples. To account for possible collector bias (e.g., different quantities of sample collection; Supplementary Figure 2), the comparative visualizations of compositional taxonomic groups were plotted separately for Dallas Zoological Gardens, Jacksonville Zoo and Gardens, and for samples collected from free-ranging elephants.

#### Extended genomic analysis and prospective applications

We assessed whether the data generated by the fecal DNA card collection and library approach presented in this study may allow for genome-level analysis. We evaluated the quality of our data by means of a phylogeographic analysis pipeline that was developed for low-coverage shotgun sequencing data (Yao et al., 2020). This pipeline uses genotype likelihoods rather than called genotypes to estimate single nucleotide polymorphisms (SNPs) in ANGSD (Korneliussen et al., 2014). Using a principal component analysis (PCA) in PCAngsd (Meisner & Albrechtsen, 2018), we determined whether the genomic data obtained from fecal cards cluster with high-quality genomes originating from the same geographic location. To generate higher coverage genomic data for fecal samples, we re-sequenced genomic libraries for three fecal card samples: SAA10 from Addo Elephant National Park, SAT04 from Tembe Elephant Park, and WNP01 from the Knysna forest. These samples represented a range of DNA quality and quantity (both in DNA concentration and percentage endogenous DNA; Supplementary Table 2 and Supplementary Table 5). Re-sequenced libraries were independently aligned to the reference genome, we then merged and deduplicated DNA alignments for each individual using SAMtools and compared these three samples to 24 geographically referenced samples (Dataset 3), resulting in a total of 27 samples. We were primarily interested in demonstrating the utility of approach by assessing clustering patterns in the PCA, and thus did not focus on inter- and intra-population differences in genetic differentiation and composition. Following Li et al. (2020), our phylogeographic approach relied on ANGSD to filter and compile a SNP dataset; we only retained SNPs that were present in at least half (14) of the 27 individuals and which had a p-value of 0.01 or less (indicative of the statistical likelihood of the position being a variable site). A total of 3,112,723,441 sites were analyzed, and 29,116,747 sites were retained after ANGSD filtering, and 12,715,208 sites were used for subsequent PCA analysis. We used PCAngsd to perform a PCA analysis, and R to visualize the clustering patterns (source code available at github.com/adeflamingh). To investigate whether small population size and inbreeding might explain the PCA clustering patterns, specifically the clustering patterns observed for the elephants from Gorongosa (see Discussion), we estimated genome-wide heterozygosity (GWH) in ANGSD as the proportion of heterozygous genotypes (analogous to theta-based estimates). Estimates of GWH have been used as a proxy for inbreeding, where inbreeding increases the homozygosity across the genome of an individual (Hansson and Westerberg, 2002; Balloux et al., 2004). WNP01 was excluded from the GWH analysis to remove low genome coverage-associated bias, and we included five-hundred million sites (“nSites = 500,000,000”) in the GWH calculation for all other individuals. In addition to the phylogeographic analysis, we also used the Rx method developed by de Flamingh et al. (2020) to estimate the biological sex of the re-sequenced fecal card sample with the lowest coverage (WNP01). We chose to estimate the sex of the sample with the lowest coverage to set a lower-bound for sequencing coverage needed to successfully estimate sex, and we did not estimate the sex of other individuals as we were only interested in demonstrating the applicability of our method, rather than investigating sex-associated genomic patterns across the datasets.

## Results

### Method Validation (Dataset 1)

To determine and quantify the effectiveness of the card collection and library building approaches that are part of our molecular pipeline, we assessed DNA concentration and endogenous elephant DNA content using repeated samples that were collected across a time interval (Dataset 1; Figure 1). The concentration of the extracted DNA was examined using broad-range (BR) Qubit quantitation and ranged from undetectable to 16 ng/μl (Supplementary Table 4). There was a significant decrease in DNA concentration as time after defecation increased (rep. measure ANOVA p = 0.0052). Consistent with the decrease in DNA concentration, the percentage endogenous DNA (Figure 1C) and the mitogenome coverage (Figure 1D) also decreased as time after defecation increased. For some samples, the DNA concentration was below the level detectable by BR Qubit quantification. However, mitochondrial genomes could be reconstructed for all sequenced samples even for those that had DNA concentrations that were too low to be quantified with BR Qubit quantitation (<1ng/uL for our samples), suggesting that DNA was still preserved in the collection card even when DNA quantitation failed. This was supported by DNA fragment analysis that showed DNA fragments of variable length were present in the genomic libraries (see Supplementary Figure 6 for an example of a DNA fragment distribution curve for one of the samples that had unquantifiable DNA). We recommend that samples be evaluated using DNA fragment analysis prior to sequencing to verify that DNA is present in the genomic library, or alternatively, using reagents with higher sensitivity (e.g., Qubit dsDNA high-sensitivity quantitation) may allow for quantitation of genomic libraries that have very low DNA concentrations (<1n/uL). More importantly, even samples collected 72hrs after defecation (the longest collection time examined) contained enough endogenous DNA to allow for high-coverage complete mitogenome reconstruction (Supplementary Table 4). For example, the complete mitogenome for the repeat 3 sample from “Jenny” could be reconstructed to 151X coverage.

Metagenomic classification of the reads in Dataset 1 into taxonomic units revealed that the composition of the samples changed as time after defecation increased for elephants from the Dallas Zoological Gardens (Supplementary Figure 4) and from the Jacksonville Zoo and Gardens (Supplementary Figure 5); in general, the proportional contribution of bacterial DNA increased, and other classified taxonomic groups (except fungi) decreased as the dung sample aged. There was also an increase in number of reads that could not be classified as time after defecation increased (Supplementary Figure 4 and Supplementary Figure 5; dark grey). The proportional contribution of endogenous elephant DNA was consistently higher in the samples that were collected from fresh dung (Rep 0) and decreased with sample age. Taxonomic classification showed that most samples contained reads that originated from bacteria, endogenous elephant DNA, arthropods, other eukaryotes, and viruses.

### Reproducibility in the field (Dataset 2)

From free-ranging elephants, collectors targeted fresh dung, and placed on the cards a larger volume of fecal matter than was collected for the zoo elephants (Supplementary Figure 2). The DNA concentration and percentage of endogenous elephant DNA did not differ significantly between fresh zoo samples (Rep 0) and samples from free-ranging elephants (Supplementary Table 5; Welch t-test for concentration, p=0.34; Welch t-test for endogenous content, p=0.87). Although the percentage of endogenous DNA did not differ significantly between samples with different amounts of fecal matter (Supplementary Figure 2), we recommend that at least 80% of each collection surface should be covered by fecal matter to maximize the possibility of obtaining and preserving elephant DNA (see Supplementary File 2: Collection protocol). DNA concentration for fecal samples from zoo elephants that were collected 24hrs, 48hrs and 72hrs after defecation (Rep 1-3) were significantly lower (Welch t-test for Rep 1 is p=0.032; Rep 2 is p=0.01 and Rep 3 is p=0.014) than the DNA concentration for samples from free-ranging individuals.

For all of the free-ranging individuals comprising Dataset 2, we were a able to reconstruct complete mitochondrial genomes (≥98% breadth of coverage), with a depth of coverage ranging from 6.9-156.6X-fold (average depth = 80X-fold) (Supplementary Table 2). We were also able to generate low-coverage genomic data for partial nuclear genomes, ranging from 3.97-35.98% of the breadth of the nuclear genome at a coverage depth of 0.01-0.60X-fold (Supplementary Table 2). Based on the proportion of reads aligning to the African elephant reference genome, samples in Dataset 2 contained an average of 12.41% (5.54-21.65%) endogenous elephant DNA. Metagenomic analysis found that on average 21.42% (0.02-34.46%) of classified reads were assigned to the genus *Loxodonta* (Supplementary Table 6).

### Nuclear genome alignment statistics for re-sequenced libraries

Three genomic libraries (SAA10, SAT04 and WNP01) were submitted for an additional round of sequencing, and alignments of the independent libraries were merged for each sample (Supplementary Table 2). For SAA10, nuclear genome coverage increased from 0.36X representing 17.85% of the breadth of the genome, to 5.8X representing 93.1% of the genome. For SAT04, nuclear genome coverage increased from 0.45X representing 30.56% of the genome, to 4.2X representing 90% of the genome. For WNP01, nuclear genome coverage increased from 0.2X representing 4.01% of the genome to 2.4X representing 14.96% of the genome. There was also a corresponding increase in coverage observed for the mitogenome, with SAA10, SAT04 and WNP01 respectively having their complete mitogenomes covered by 1248X, 668X and 193X reads.

### Extended genomic analysis and prospective applications (Dataset 1, 2 and 3)

#### Metagenomics

Metagenomic analysis revealed that fecal samples from free-ranging elephants contained reads that originated from the same main classification groups (*Loxodonta*, green plants, fungi, arthropods, bacteria, viruses, archaea) that were detected in zoo individuals. Zoo and free-ranging elephant samples did not contain reads of human origin except for a single zoo sample (Thandi, Rep 2) and a single free-ranging elephant sample (SAA01), both of which had ≤ 0.01% of total reads assigned as originating from *Homo sapiens*. The average number of classified reads assigned to *Loxodonta* was higher for free-ranging (21.42%) than zoo (7.9%) individuals, and the relative contribution of bacteria was lower in free-ranging individuals (Figure 2, brown portion) while other taxonomic groups such as arthropods (Figure 2, black portion) were more abundant in free-ranging than zoo samples.

**Figure 2:**
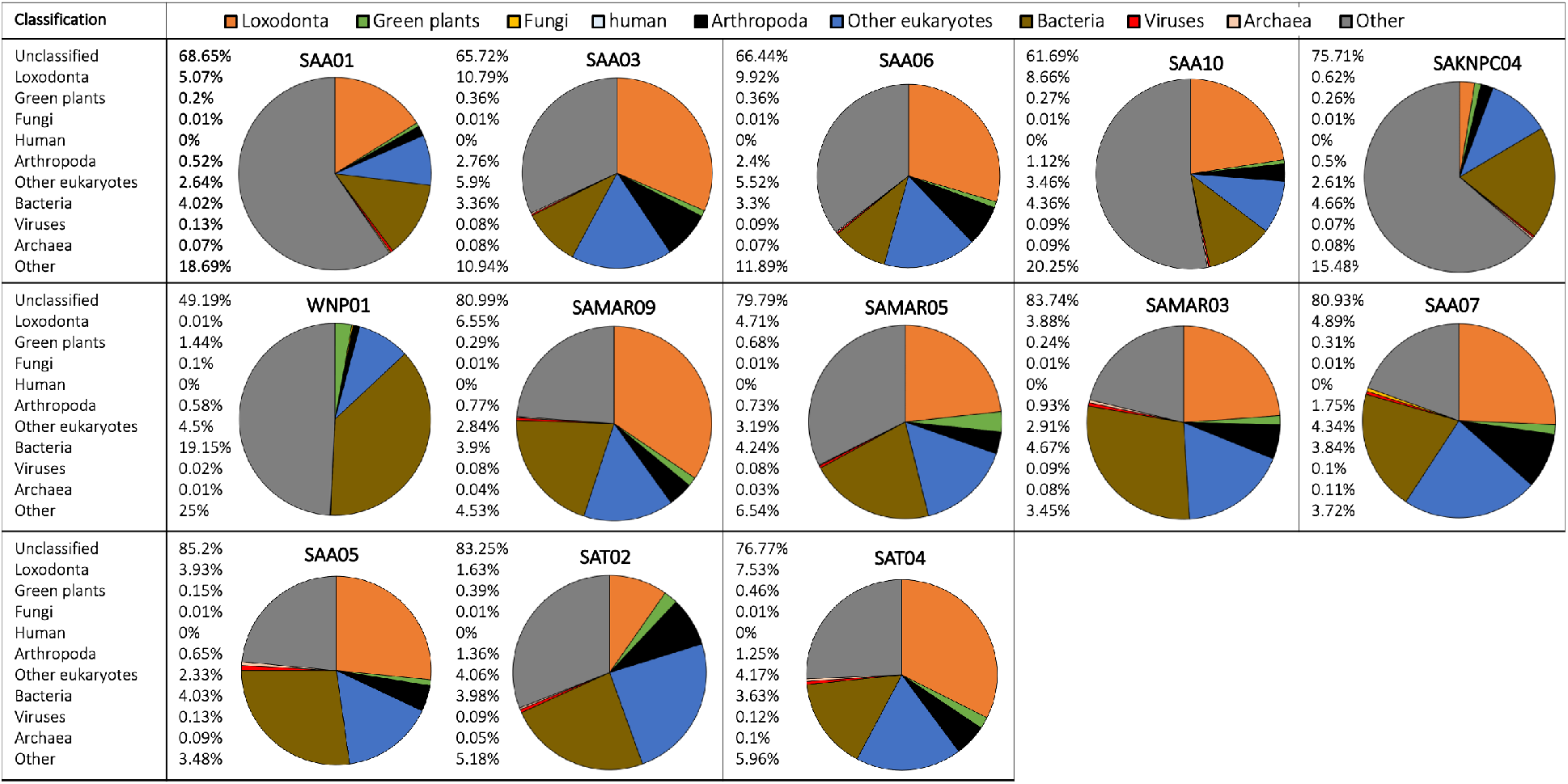
Metagenomic classification of fecal DNA samples from 13 free-ranging elephants from South Africa. Compositional taxonomic contributions of classified reads are summarized as a pie chart for each individual (unclassified reads are not shown). Taxonomic groups have been color coded and corresponding colors are indicated in the legend in the top row of the figure. Taxonomic classification showed that most samples contained reads that originated from bacteria, endogenous elephant DNA, green plants, arthropods, and other eukaryotes.

Most samples (zoo and free-ranging) contained reads that originated from arthropods. For example, samples from fresh dung (Rep 0) for “Jenny” and “Mlilo” from Dallas Zoological Gardens contained reads originating from Arthropoda (Supplementary Figure 4). For these two samples, arthropods respectively contributed 0.68% and 0.89% of all reads (including unclassified reads), or 3.2% and 4.1% of classified reads. Of these classified reads, a substantial proportion (45% and 42.7% of reads classified as Arthropoda) was identified as belonging to the lepidopteran clade Ditrysia. Similar patterns and taxon identifications were observed for samples collected from free-ranging individuals (e.g., SAA03, SAA06 and SAT02 contained a substantial proportion of reads originating from Ditrysia).

For some individuals a proportion of the reads originated from Archaea. For example, “Thandi” from the Jacksonville Zoo and Gardens (Supplementary Figure 5) had a substantial portion or reads assigned to the genus *Methanobrevibacter*, a dominant gut-associated archaeon (Hansen et al 2011). There were relatively few reads classified as originating from Fungi across all samples (zoo and free-ranging), with the highest proportion of fungal reads present in the samples for which the most time had passed since deposition (e.g., Rep 3 samples for Ali and Thandi; Supplementary Figure 5). Metagenomic analysis also identified reads originating from viruses, e.g., multiple samples (SAA05, SAT04; Figure 2) contained contributions from Caudovirales.

#### Principal components analysis (PCA) and genetic sex

A PCA using nuclear genome-wide SNPs revealed geographical partitioning of the elephants (Supplementary Figure 1). Elephants for which fecal card samples were collected from Addo Elephant National Park and Tembe Elephant Park in South Africa grouped most closely with geo-referenced elephants (Dataset 3) that were also from South Africa (e.g., Kruger) (Figure 3B; cluster at top left of panel). The Knysna elephant from South Africa (fecal card sample WNP01) did not cluster with any of the geo-referenced elephants when considering PC1 (Figure 3B, Supplementary Figure 7A). However, all three elephants from South Africa from which DNA was collected using fecal cards did cluster together when PC2 and PC3 were considered (Supplemental Figure 7B). Each of the principal components contributed very little to the overall genomic variation observed (PC1: 2.8%, PC2: 1.78%, PC3: 1.67%). Elephants from Gorongosa National Park, Mozambique were separated by PC1 and PC2 into three clusters (Figure 3A, blue symbols). The partitioning elephants from Gorongosa National Park did not correspond to genome-wide heterozygosity (Supplementary Figure 8), and the overall clustering pattern for all elephants was not concomitant with genome sequence coverage (Supplementary Figure 9; all clusters included a range of coverages). Finally, we examined whether the biological sex of the elephants could be determined using DNA sequencing reads from fecal cards. We examined the elephant with the lowest genome coverage of the three South African elephants, the Knysna elephant (WNP01), and were able to identify this elephant as a female (Rx: 0.8274169, p< 0.05) using the approach of de Flamingh et al. (2020).

**Figure 3:**
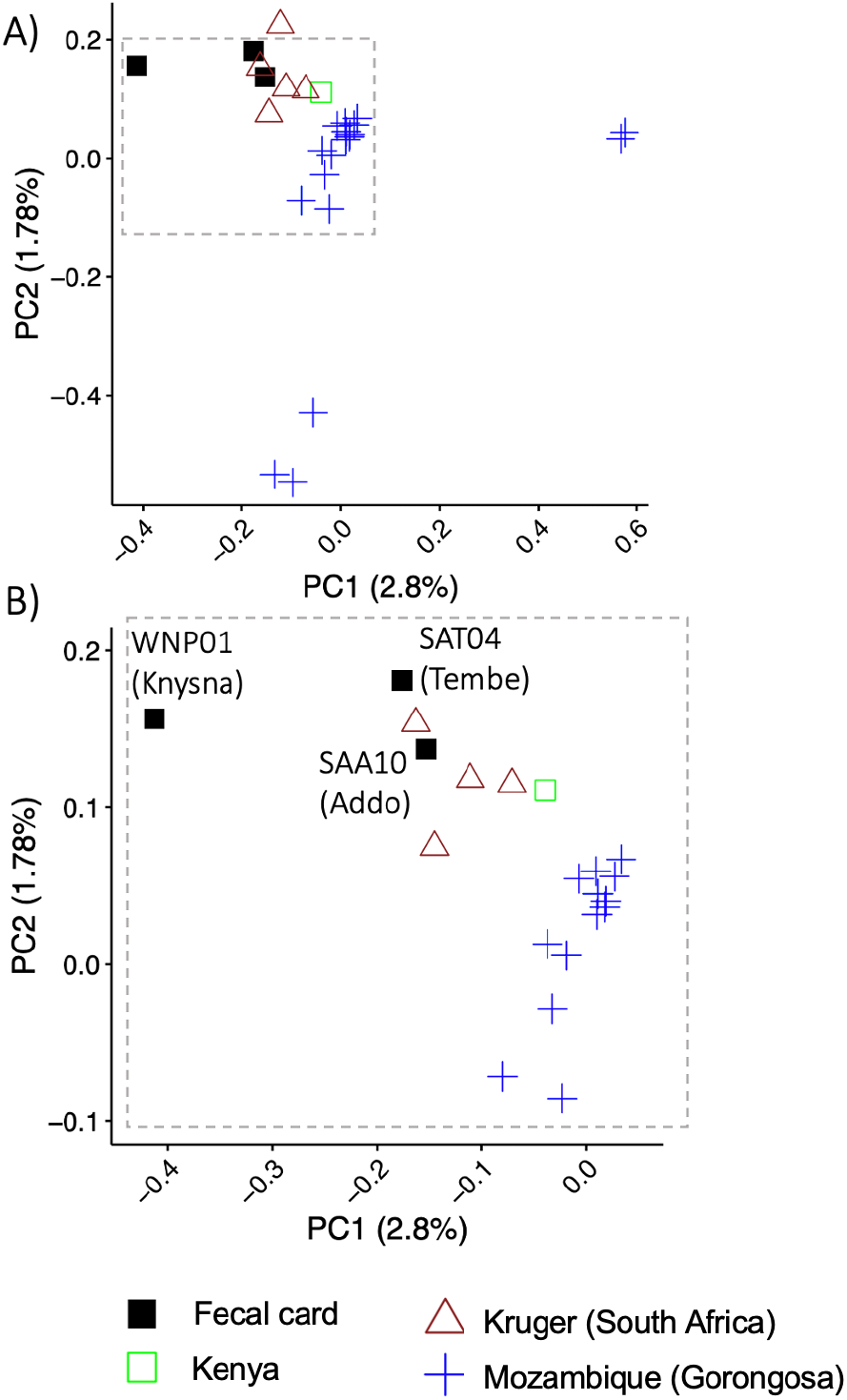
Assessment of nuclear genetic variation for free-ranging elephants from South Africa, Mozambique and Kenya. Principal component analysis using nuclear genome-wide SNPs (single nucleotide polymorphisms) supported the geographic partitioning of elephants (n=27) in an analysis combining sequences of DNA from elephant dung collected on DNA preservation cards (“fecal card”) in South Africa with sequences of high-quality DNA from geo-referenced elephant samples from Kruger (South Africa), Mozambique and Kenya. Elephant DNA from dung collected on DNA preservation cards in Addo Elephant National Park and Tembe Elephant Park group with geo-referenced individuals from Kruger National Park in South Africa that were sequenced using high quality DNA. The Knysna elephant fecal sample (WNP01) does not cluster with the other South African elephants. Panel A shows the clustering pattern for all 27 elephants using PC1 and PC2; individuals from Gorongoza do not overlap with elephants from other locations, while the single Kenya elephant is outside the range of values for South African elephants in PC1. Panel B shows an enlargement of the top left cluster of individuals in panel A. Symbols and colors represent different geographic localities (legend below the panel).

## Discussion

We present a molecular pipeline that incorporates a simplified and accessible DNA sample collection and extraction protocol that uses a DNA preservation card which does not require refrigeration after sample collection, and is easy to transport and store in the field and in the laboratory. Other fecal DNA sample collection methods rely on refrigeration and/or chemical preservation (Murphy et al., 2002; Nsubuga et al., 2004). Access to refrigeration is often not feasible due to mode of transport (e.g., itinerant and/or trekked field expeditions) or because of limited access to electricity in undeveloped/rural areas. Previous means of chemical preservation of elephant fecal samples for DNA analyses have involved chemicals that may be cumbersome to handle, may be dangerous, and may require refrigeration, e.g., ethanol or preservation buffers Wasser et al., 2004; Gobush et al., 2008; Gray et al., 2011). In addition to these risks, the international transport or shipment of chemicals (e.g., ethanol) is often regulated and sometimes prohibited. Such methods of DNA sample collection may therefore hinder research projects. The use of fecal DNA preservation cards avoids these issues and provides an easy and accessible way for in-field researchers and conservation practitioners to collect and effortlessly transport many samples at once.

Our approach tested and allowed for DNA analysis of fecal samples that were up to 72hrs (3 days) old, meaning that samples could be opportunistically collected in the field even days after animals have left an area (Figure 1). We were able to generate complete mitogenomes for all shotgun-sequenced zoo individual samples collected 72hrs after defecation (>90% of the mitogenome with an average coverage of 55.7X for the samples). However, samples that were collected shortly after defecation more consistently resulted in high DNA concentration, high mitogenome coverage, and high endogenous target-species DNA content. To maximize the possibility of successful DNA collection, sample collectors should ensure that samples are completely dry before long-term storage, as DNA is prone to degradation if samples are not completely desiccated (Murphy et al., 2002; Nsubuga et al., 2004). We recommend storing DNA preservation cards with a desiccant (Supplementary File 2 – Step 5). The preservation card collection protocol follows 5 easy steps, and no specialized training is required, therefore, samples may also be collected by persons who do not have formal scientific research training. In addition, the DNA preservation cards used in our approach are relatively inexpensive (at the time of this study the cost was less than US$5.00 per card with two collection surfaces for each bolus of dung).

The pipeline presented in this study does not require host targeted DNA enrichment (Chiou and Bergey, 2015) for the sequencing of complete mitochondrial genomes and low-coverage nuclear genomes. On average our method resulted in 10-20% endogenous host-specific DNA (based on bioinformatic and metagenomic statistics; Supplementary Table 2, Supplementary File 4). We show that library re-sequencing can produce datasets that represent >90% of the complete nuclear genome of the target species, with coverages up to 6X and potentially higher. However, the resulting coverage of re-sequenced libraries is dependent on the diversity of available DNA template in the original library (Daley and Smith, 2014), as limited DNA template diversity will result in replicate sequencing of the same DNA molecule, resulting in PCR duplicate reads rather than unique reads that contribute to the breadth (%) and depth (X-fold) of genome coverage. For example, initial sequencing of WNP01 resulted in a comparatively smaller percentage of nuclear genome coverage (4% compared to ~17% in the other two resequenced individuals), and the re-sequencing of this library was less productive (~15% of the nuclear genome was recovered, compared to >90% for the other two re-sequenced libraries), likely due to limited DNA template molecule diversity. We therefore encourage researchers to conduct an initial sequencing screen to calculate genome coverage statistics that can be used to inform the choice of libraries to re-sequence. Initial screening of sequencing data can be conducted using software that predicts library complexity and prospective genome coverage, e.g., the program “preseq” (Daley et al., 2014).

Sequencing efforts of libraries generated through our approach will need to be increased to obtain target-species genome coverages similar to that of other methods that use bait or other enrichment protocols. However, the time, effort, and reagent cost associated with complex DNA- and RNA-bait design and laboratory enrichment may offset the potential cost of increased sequencing, especially as the costs of sequencing may continue to fall. In addition, our approach may outperform other methods in that it also allows for host-*associated* DNA to be investigated (e.g., DNA that is not from the host genome, but is extracted as part of the fecal sample, for example, fecal microbiome data).

The ability to generate nuclear genome-wide data from non-invasively collected fecal samples can open up new possibilities for genomic analysis. We demonstrated the utility and potential application of our approach by generating and analyzing data for African savanna elephants. Our approach may also be directly applicable to other taxa. We showed that the nuclear SNP dataset generated for non-invasive fecal DNA from free-ranging elephants is sufficient for complex molecular and bioinformatic analyses. For example, our analysis indicated that the fecal card samples grouped with other individuals from South Africa (e.g., individuals from Kruger), supporting the expectation of clustering of individuals that are geographically close together, as reported by earlier studies that used nuclear DNA microsatellites (Ishida et al., 2011; Wasser et al., 2015). However, there are unresolved patterns (e.g., the positioning of the Mozambique elephants, Figure 3) that would need to be further explored through a comparison of Dataset 3 to continent-wide genomic data, although beyond the scope of the current study with its primary focus is on reporting a method for fecal DNA sample analysis. The possibility of generating genome-wide data from fecal samples may also shed light on other broad-scale genomic questions, for example, the evolutionary history of elephants and other species, and estimation of the age of inter-species hybridization zones (Tonzo et al., 2020; Bonnald et al., 2021).

In the PCA, the geo-referenced elephants from Gorongosa National Park in Mozambique were separated into three clusters when considering PC1 and PC2 (Figure 3). These elephants were part of a study that investigated tusklessness as a X chromosome–linked dominant, male-lethal trait(Campbell-Staton et al., 2021). The elephant population in Gorongosa has fluctuated drastically during the last three decades, with a >95% decrease in population size as a consequence of civil war (1977-1992), and limited recovery of the population since the end of the civil war (Stalmans et al., 2019; Campbell-Staton et al., 2021). In addition, the population was augmented through the translocation of 6 elephant bulls from Kruger National Park in 2008 (Stalmans et al., 2019), which may have affected clustering in the PCA. We investigated the possibility that the PCA clustering pattern of Gorongosa elephant genetic variation may be a consequence of small population size and associated inbreeding (Allendorf et al., 2013) by estimating genome-wide heterozygosity (GWH) for Gorongosa and other elephants in this study (except WPN01 that was excluded due to low genome coverage). The GWH was not associated with the clustering patterns observed for Gorongosa elephants (Supplementary Figure 8), nor by the tusk status of these elephants (Campbell-Staton et al., 2021). Future research that compares the nuclear genomes of these elephants to geo-referenced nuclear genomes from elephants across different regions in Africa on a continental scale, including higher representation of regions surrounding Mozambique and more individuals from Kruger National Park, might shed light on the observed pattern of genetic variation.

In addition to broad-scale conservation genomic questions (e.g., phylogeography and genetic structuring), our approach may enable the study of rare or elusive species, or for species or populations where low animal density or small population size make traditional methods of sample collection (e.g., trapping/immobilization) difficult or impossible. For example, using the approach developed by de Flamingh et al. (2020), we estimated the genomic sex of the only remaining Knysna elephant in Wilderness National Park (WNP; Moolman, de Morney, et al., 2019). This elephant which lives in the Afromontane Knysna Forest of South Africa is the sole survivor of a population that once comprised thousands of elephants, and there is ongoing discussion on management actions (e.g., reintroduction of other elephants; (Moolman, de Morney, et al., 2019; Moolman, Ferreira, et al., 2019; Patterson, 2012)). Consistent with photographic evidence (Moolman et al., 2019a), we confirmed that this is a genetically a female. Stakeholders should take this into consideration when developing management plans. This individual does not group with other georeferenced elephant samples. This may represent a unique relic genomic signature, but this clustering pattern could also be driven by low genome coverage rather than genetic variation.

Unlike previous studies that sought to generate nuclear genome data from non-invasive samples (Perry et al., 2010; Chiou and Bergey, 2015; Snyder-Mackler et al., 2016; Taylor et al., 2021), our DNA extraction and genomic library construction approaches did not target host-specific DNA. Non-target enriched shotgun sequencing data have been used to reconstruct mitochondrial genomes (Bon et al., 2012; Srivathsan et al., 2019; Ang et al., 2020), but low endogenous/target DNA content have so far precluded the generation of nuclear genome-wide datasets without targeting host-specific DNA (Perry et al., 2010). To overcome low endogenous DNA in non-invasive samples, researchers have used DNA and RNA baits and methylation patterns to enrich genomic libraries for host species DNA prior to sequencing (Perry et al., 2010; Chiou and Bergey, 2015; Snyder-Mackler et al., 2016). Bait-based and other enrichment protocols are expensive, labor-intensive and time consuming, and may result in capture-biases where portions of the genome may be overrepresented in the sequencing pool after enrichment (George et al., 2011; Chiou and Bergey, 2015). Our molecular pipeline precluded such obstacles by shotgun-sequencing total genomic DNA using standard DNA extraction and library construction protocols, and allowed us to capitalize on the availability of concomitant sources of DNA that form part of the collected fecal sample. Sequencing host-associated DNA decreases the relative contribution of endogenous host-specific DNA when samples are not enriched, but conversely this allows for the study of complementary aspects that can inform the conservation and management of species. Because parasites and pathogens may be identified in fecal samples (Srivathsan et al., 2019), our approach has the potential for monitoring individual and population health, which may be especially beneficial for rare, elusive, and endangered species whose health cannot be assessed using traditional approaches (e.g., blood sample screening). The ability to assess and monitor microbiome communities, and therefore individual and population health, may inform species conservation and management (Kophamel et al., 2022). For example, our metagenomic analysis of host-associated DNA showed that multiple samples contained contributions from Caudovirales, an order of bacteriophages that has been associated with gut health (Lepage et al., 2008; Minot et al., 2011; Norman et al., 2015). Fecal samples can also be used to examine intestinal parasite presence and abundance (e.g., Srivathsan et al., 2019), and fecal DNA analysis is an effective and reliable method for studying parasite infections (da Silva et al., 1999; Blessmann et al., 2003; Yoshikawa et al., 2011). In our free-ranging elephants (Supplementary File 4) reads were identified as originating from the phylum *Platyhelminthes*, which includes flatworms that are predominantly parasitic (Park et al., 2007).

The metagenomic analysis that we applied in this study showcases the potential utility of our approach for addressing complementary questions based on host-associated DNA that can also inform species conservation, for example, through pathogen and parasite identification. The pipeline presented here may expand the application of genomic techniques to conservation science.

## Supporting information

Supplementary File 2 Collection protocol

Supplementary File 4 K2 output

## Acknowledgements

For funding, we thank the International Fund for Animal Welfare; the Conservation Ecology Research Unit (CERU) of the University of Pretoria; and the US Fish and Wildlife Service African Elephant Conservation Fund, grant F22AP01215-00. Alida de Flamingh was supported by funding from the University of Illinois at Urbana-Champaign. Patrícia Pečnerová received funding from the European Union’s Horizon 2020 research and innovation programme under the <u>Marie Skłodowska-Curie</u> grant agreement No 892446.For samples from free-ranging elephants, we thank CERU. For samples from elephants in zoos, we thank Lara Metrione and Linda Penfold of the South-East Zoo Alliance for Reproduction and Conservation; Dianna Lydick, Lora Baumhardt and the Dallas Zoo; and Fatima Ramis, Corey Neatrou and the Jacksonville Zoo and Gardens.

## Conflict of Interest statement

The authors declare no conflicts of interest with respect to the content, authorship, and/or publication of this article.

## Data Accessibility and Benefit-Sharing

Access to published genomic datasets are listed in Supplementary Table 1. Genomic data for the zoo and free-ranging elephants is available on the Short-Read Archive under bioproject number (XXXXXXX), metagenomic results is available on DRYAD (XXXXXX).

## Author Contribution statement

AdF, RJvA, RSM and ALR conceptualized and designed this study; RJvA provided free-ranging elephant samples; HRS, PCP and YI provided access to high quality geographically referenced data; AdF and SV carried out the molecular analysis of fecal samples; AdF and YI carried out bioinformatic and statistical analyses; AdF, RJvA, RSM and ALR interpreted the data and results; all authors contributed to and approved the final version of this manuscript.

## Notes

### Competing Interest Statement

The authors have declared no competing interest.

