## Supplementary File 2 Collection protocol for "Non-invasive fecal DNA yields whole genome and metagenomic data for species conservation"

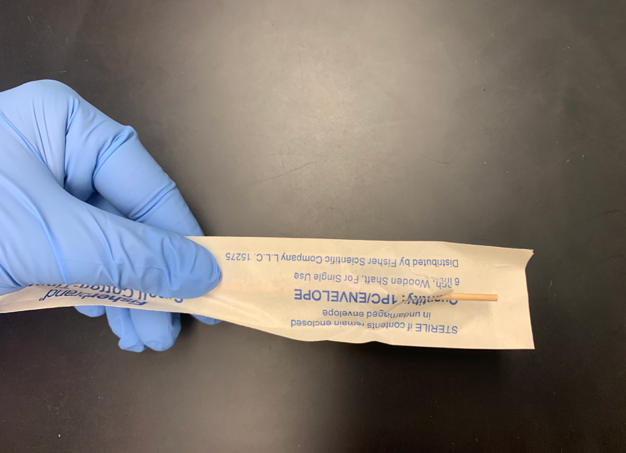
**SAMPLE COLLECTION PROTOCOL CONTINUES ON NEXT PAGE**

Alternatively, if no cotton-tip applicator is available, please **collect the sample from any location on the outside surface of the dung ball by pressing the card directly against the dung surface for 15 seconds**.

**Sample collection protocol using GeneSaver or Whatman FTA cards**

**Samples should be collected upon or shortly after defecation**. For each card, there will be 2 collection surfaces (circles – please see images below). **Please make sure to add the sample that you are collecting so that it covers both of the collection surfaces/circles in the card (step 5).** Please ensure that the entire circle is covered with fecal matter (see example, step 5). Please complete and add sample ID and GPS coordinates to the collection card AND the attached collection spreadsheet.

Step 1: Put on sterile gloves. Please use a new pair of gloves each day and for each elephant.

Step 2: Label each collection card with **individual/sample name and GPS coordinates**, in the location indicated below by red box

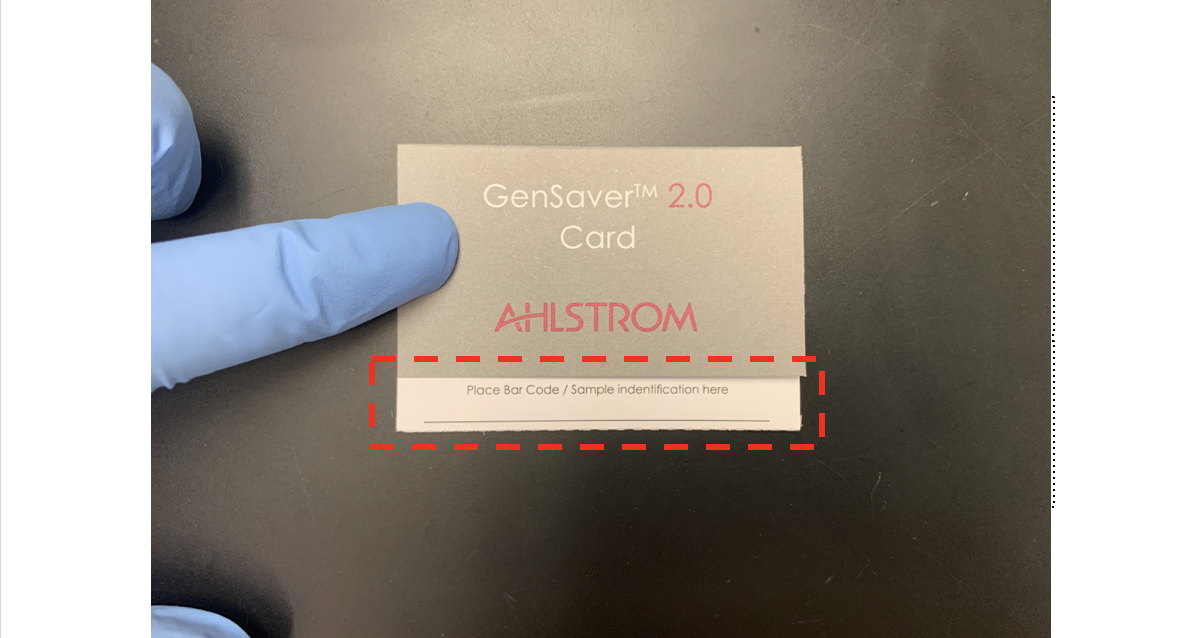

Step 3: Open the cotton-tip applicator and press the cotton-tip against the outside of the elephant dung ball. Roll the cotton-tip from side to side so that the tip is saturated with fecal material. Alternatively, if no cotton-tip applicator is available, please collect the sample from any location on the outside surface of the dung ball by pressing the card directly against the dung surface for 15 seconds.

Step 4: Open the sample collection card to expose the two circles that contain the preservative paper on which the fecal sample should be applied. Do not touch the preservative paper directly with gloves.

Step 5: Press cotton tip against the collection card on the inside of the delimited circles which contain the DNA preservative. Aim to **cover at least 80% of each circle with fecal sample** from the cotton-tip applicator (see example below). It is often the case that too little volume is collected – collectors should ensure that a sufficient layer of fecal material is present on each circle. Alternatively, if no cotton-tip applicator is available, please collect the sample from any location on the outside surface of the dung ball by pressing the card directly against the dung surface for 15 seconds. The same gloves and applicator can be used for both samples in one card, but please use a new set of gloves and applicator for each individual card and elephant.

Example of collected sample:

Step 5**: Leave card open overnight (or for at least 4h)** so that the fecal sample dries completely before closing and shipping the sample. The sample can be shipped at room temperature. Please place each sample card in the provided Ziplock bag before shipment to avoid cross-contamination. Please place one sachet of desiccant in each of the Ziplock bags.

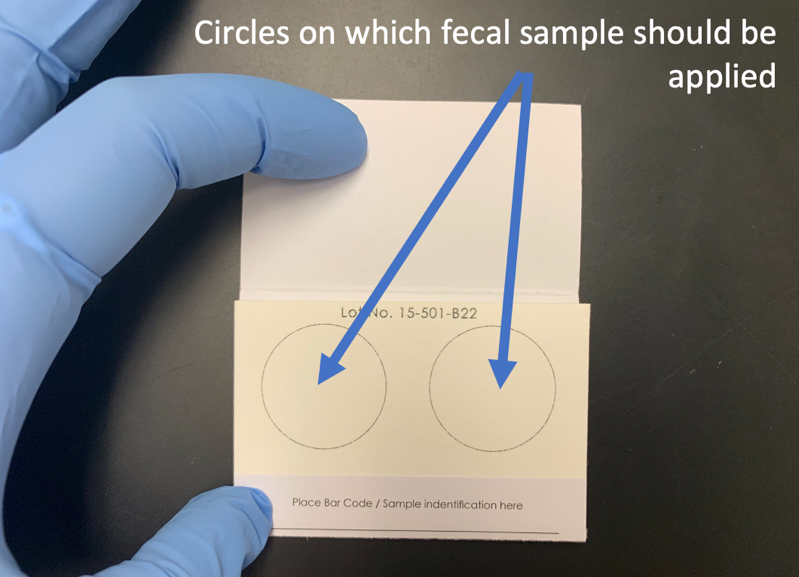

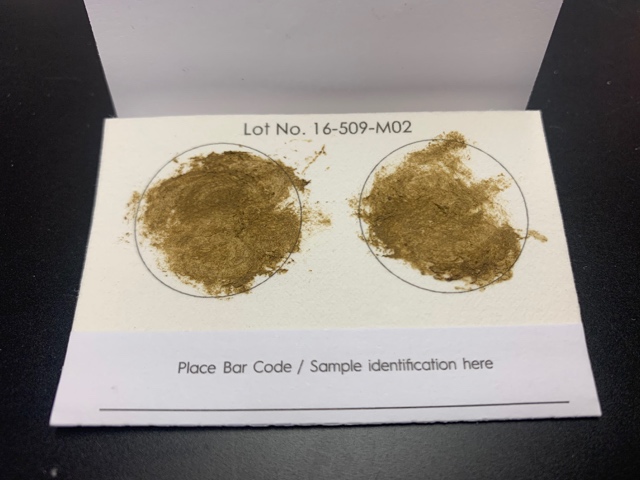

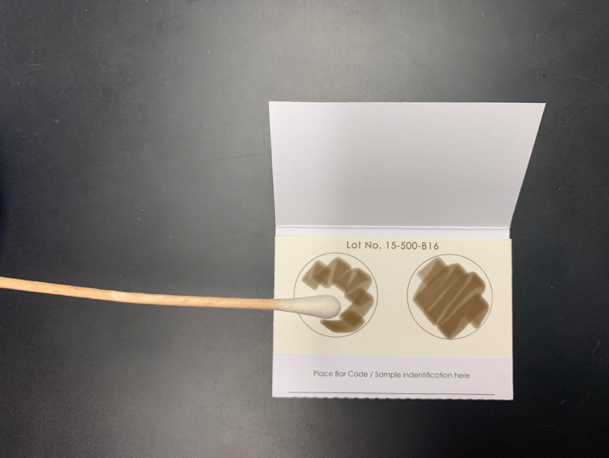

Collection Spreadsheet:

|  | Sample name: | Sample coordinates: | Date | Dung circumference | Collector | Additional notes** |
| --- | --- | --- | --- | --- | --- | --- |
| 1 |  | 1)  2) |  |  |  |  |
| 2 |  | 1)  2) |  |  |  |  |
| 3 |  | 1)  2) |  |  |  |  |
| 4 |  | 1)  2) |  |  |  |  |
| 5 |  | 1)  2) |  |  |  |  |
| 6 |  | 1)  2) |  |  |  |  |
| 7 |  | 1)  2) |  |  |  |  |
| 8 |  | 1)  2) |  |  |  |  |
| 9 |  | 1)  2) |  |  |  |  |
| 10 |  | 1)  2) |  |  |  |  |
| 11 |  | 1)  2) |  |  |  |  |
| 12 |  | 1)  2) |  |  |  |  |
| 13 |  | 1)  2) |  |  |  |  |
| 14 |  | 1)  2) |  |  |  |  |
| 15 |  | 1)  2) |  |  |  |  |
| 16 |  | 1)  2) |  |  |  |  |
| 17 |  | 1)  2) |  |  |  |  |
| 18 |  | 1)  2) |  |  |  |  |
| 19 |  | 1)  2) |  |  |  |  |
| 20 |  | 1)  2) |  |  |  |  |

** Additional notes may include, for example: landscape/habitat in which sample was collected, sex of elephant if known, and any additional comments on sample.
