## Supplementary File 3 Figures for "Non-invasive fecal DNA yields whole genome and metagenomic data for species conservation"

### Supplementary Figures – de Flamingh et al.

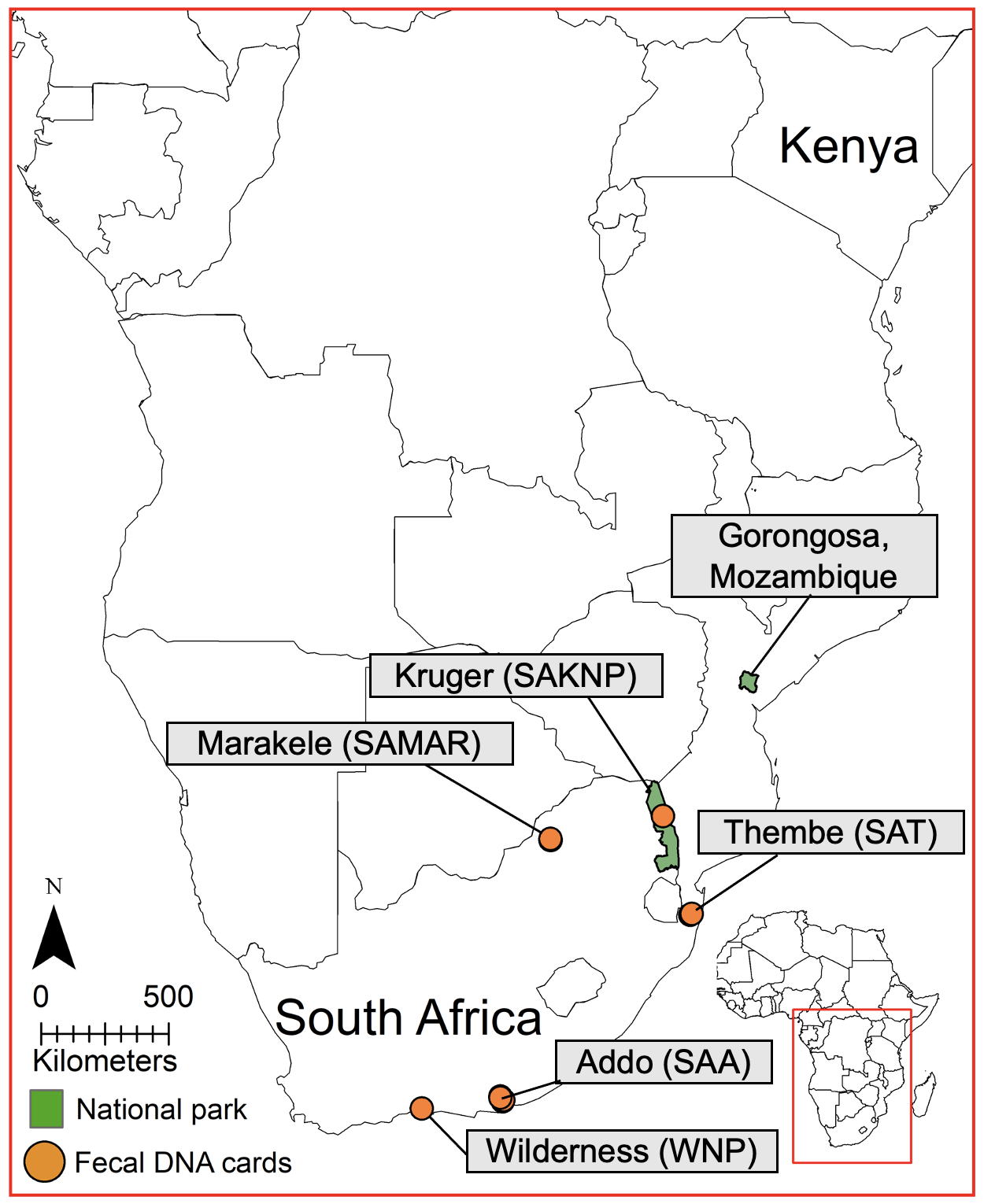

Supplementary Figure 1. Map of southern Africa showing the geographic provenance of sampled elephants. Our study comprised 3 datasets; one dataset of 24 samples from 6 elephants under human care from two zoos in the United States of America (Dataset 1), a second set of 13 wild African savanna elephants (*Loxodonta Africana*) collected from 5 geographic areas within South Africa using fecal DNA preservation cards (Dataset 2, orange circles), and a third set of reference 24 high-quality samples (Dataset 3): 5 samples from Kruger National Park, South Africa, one sample from Kenya and 18 samples from Gorongosa National Park, Mozambique. Abbreviations for geographic locations are listed in brackets following the park or area name, e.g., Wilderness National Park (WNP), and are consistent across figures and supplementary tables.

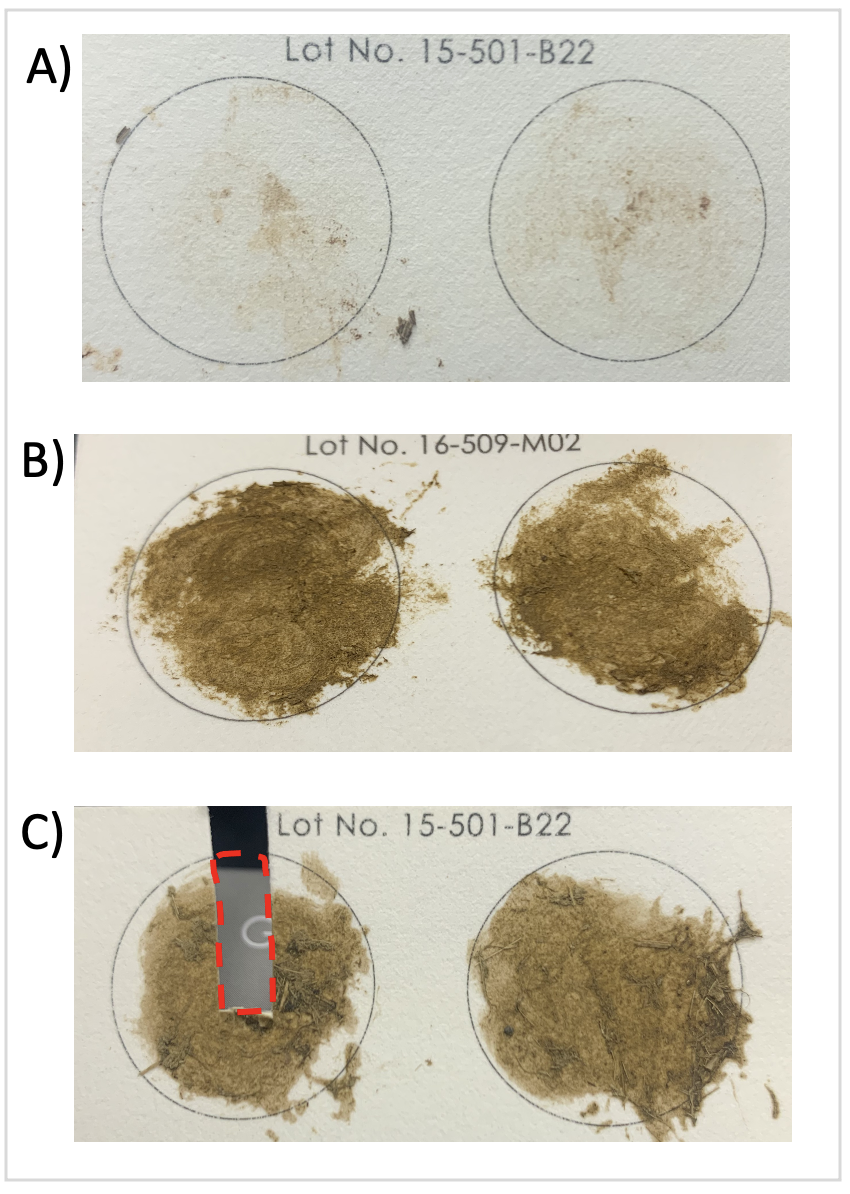

Supplementary Figure 2. Thirty-seven cards were used to collect fecal samples from zoo (A) and wild (B) elephants using a standardized collection procedure (Supplementary File 2). Sample cards from zoo individuals contained less fecal material than what was collected for wild individuals; however, extracted DNA concentration did not differ for the fecal samples between those collected in the wild and those collected fresh in zoos. (C) Approximately one-quarter of one of the two collection surfaces in each card was used as starting template for DNA extraction using the QIAamp® PowerFecal® Pro DNA Kit (Qiagen, Ann Arbor, United States).

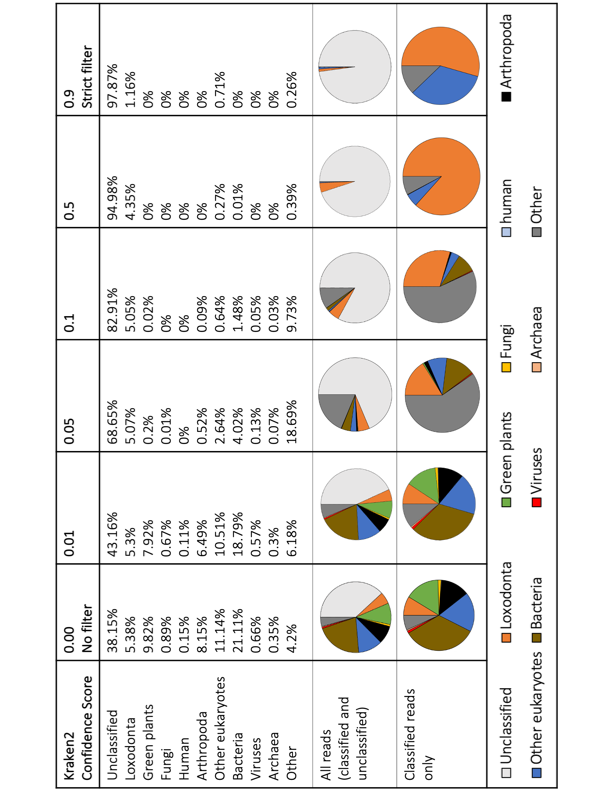

Supplementary Figure 3. A range of confidence score (CS) threshold values was assessed for metagenomic classification in Kraken2 (Wood et al., 2019). The CS threshold filtering approach is implemented in Kraken2 to allow users to specify a threshold score between 0 and 1; the classifier then adjusts labels up the classification tree until the label's score meets or exceeds that threshold (see Wood et al., 2019 for details). If a label at the root of the taxonomic tree does not have a score exceeding the threshold, the sequence is called unclassified by Kraken 2 when this threshold is applied. This approach minimizes misclassification and false positive rates associated with metagenomic data classification. We tested the classification without any confidence filtering (CS = 0.00), and with CS thresholds of 0.01, 0.05, 0.1, 0.5, 0.9 for a sample from a wild elephant (SAA01) with DNA concentration and genomic depth and breadth of reads (mapped to the African elephant genome) typical of our data (see Figure). Each column represents the results of a different classification threshold, and pie-charts show the taxonomic break-down of all reads (top chart) and classified reads only (bottom chart). Taxon identities have been color coded and are consistent with colors used in the main manuscript. We found that successful sequence classification is inversely related to the threshold stringency. We found that a filtering threshold of 0.05 is sufficient to remove false positive rates (FPR) and erroneous classification without impacting successful classification drastically (i.e., reads classified as “human” were removed from results when a classification threshold of 0.05 was applied). In addition, this analysis showed that Kraken2 is effective and accurate when assigning reads to elephant/target-species DNA as the program consistently classified ~5% of reads from this sample as “Loxodonta”, regardless of the confidence threshold specified.

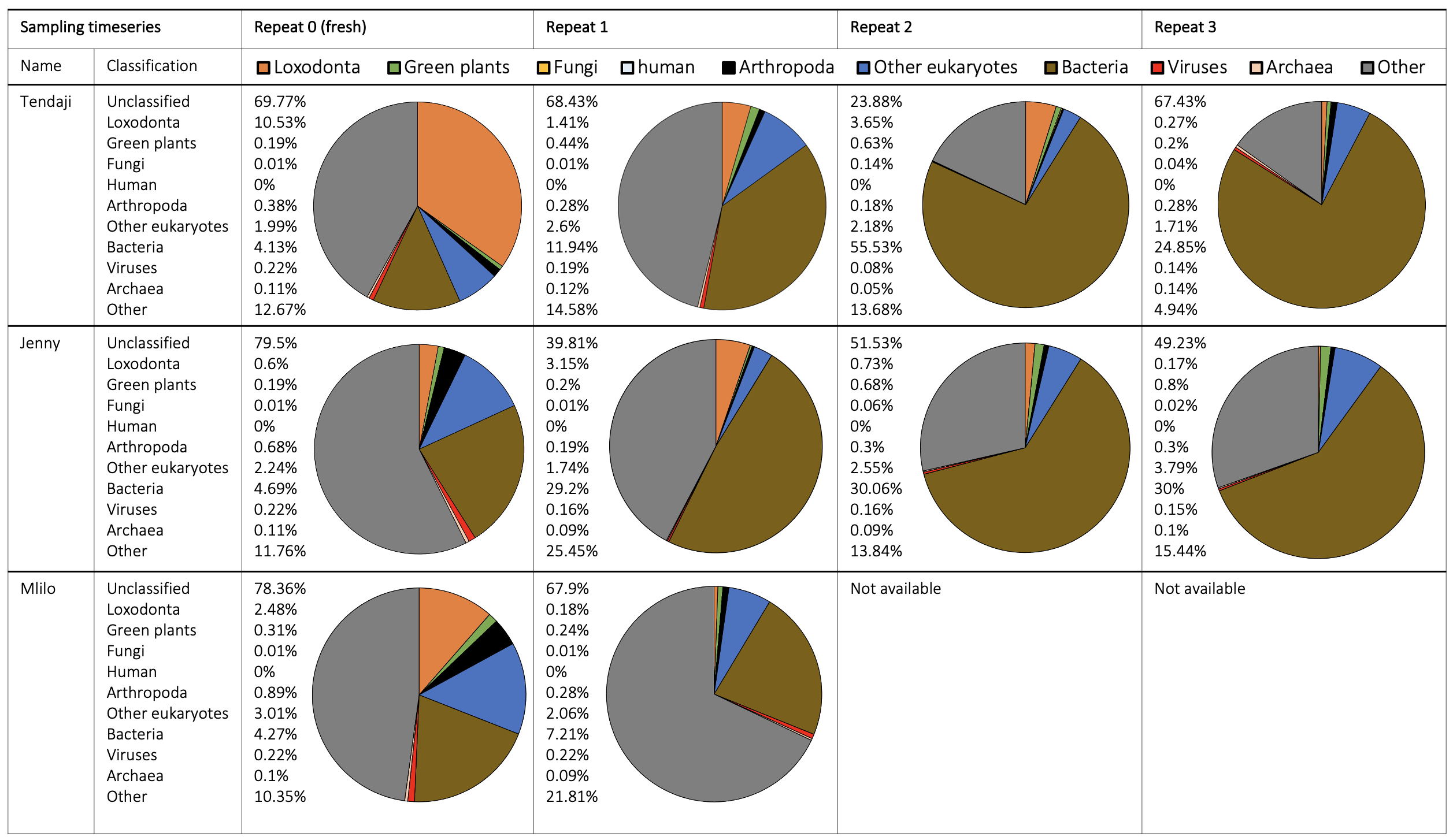

Supplementary Figure 4. Metagenomic classification of fecal DNA samples from three individuals from the Dallas Zoological Gardens (Tendaji, Jenny, Mlilo) into taxonomic units revealed that the composition of the samples changed as time after defecation increased; the proportion of bacterial DNA increased, and all other classified taxonomic groups decreased as the dung sample aged. There was also an increase in the proportion of reads that could not be classified as time after defecation increased. The proportion of endogenous elephant DNA was consistently higher in the samples that were collected from fresh dung (Replicate 0) and decreased with sample age. Taxonomic classification showed that most samples contained reads that originated from bacteria, endogenous elephant DNA, arthropods, other eukaryotes, and viruses. Pie charts show the proportional contribution of *classified* reads per taxonomic group, reads that were unclassified are not shown in the pie charts.

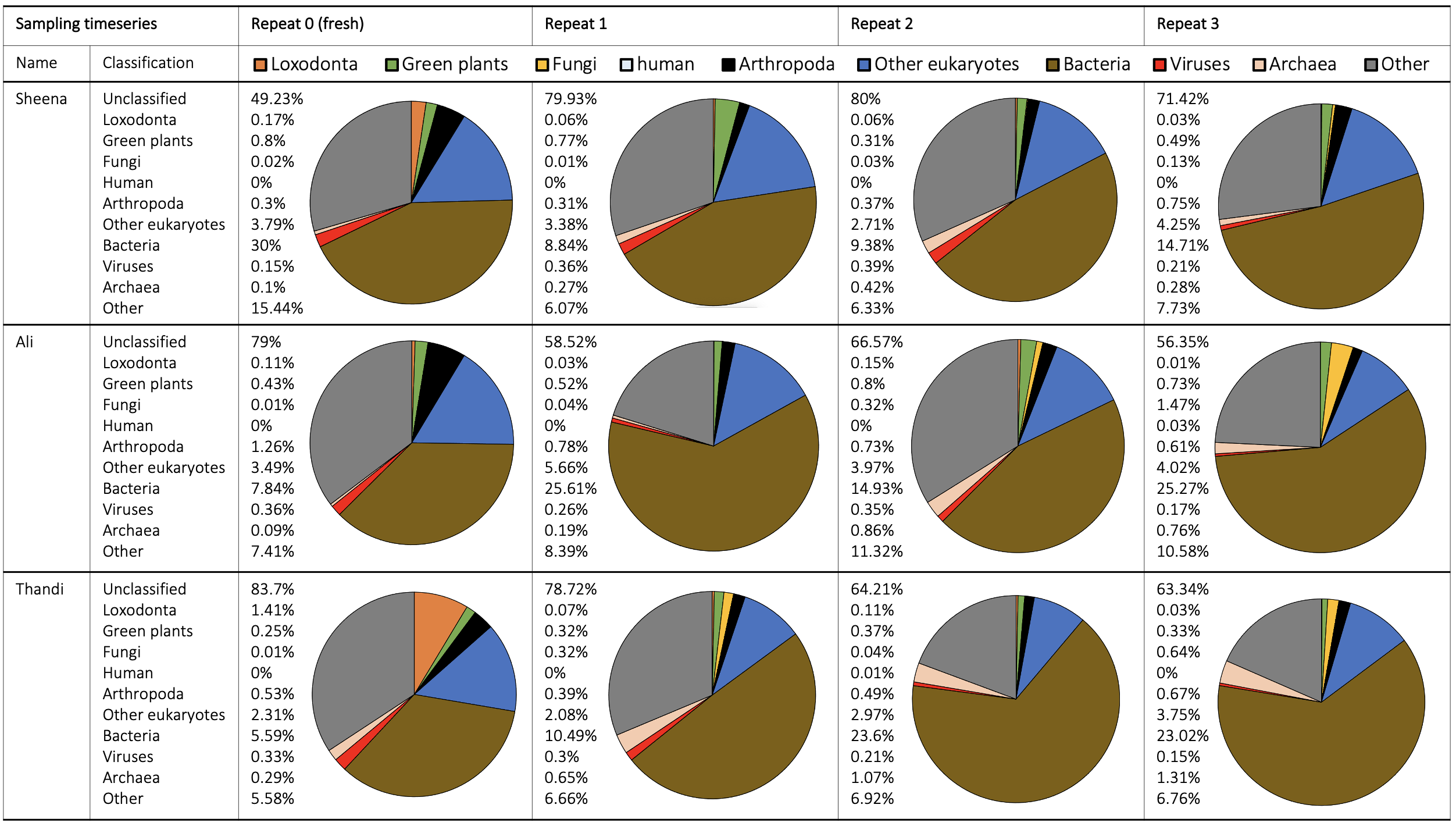

Supplementary Figure 5. Metagenomic classification of fecal DNA samples from three individuals from the Jacksonville Zoo and Gardens (Sheena, Ali, Thandi) into taxonomic units revealed that the composition of the samples changed as time after defecation increased; the proportion of bacterial DNA increased, and all other classified taxonomic groups decreased as the dung sample aged. There was also an increase in the proportion of reads that could not be classified as time after defecation increased. The proportion of endogenous elephant DNA was consistently higher in the samples that were collected from fresh dung (Replicate 0) and decreased with sample age. Taxonomic classification showed that most samples contained reads that originated from bacteria, endogenous elephant DNA, arthropods, other eukaryotes, and viruses. Pie charts show the proportional contribution of *classified* reads per taxonomic group, reads that were unclassified are not shown in the pie charts.

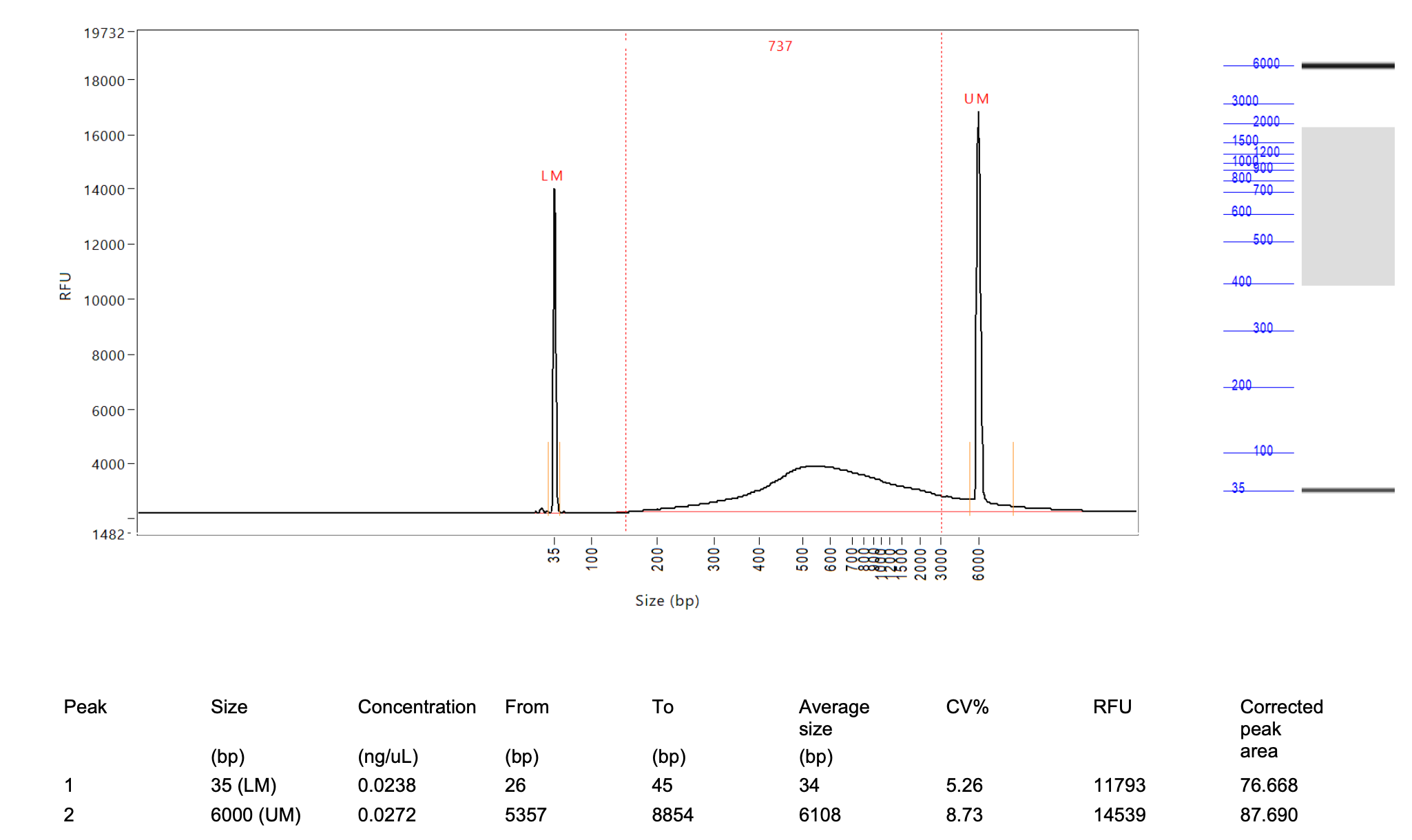

Supplementary Figure 6. DNA fragment size analysis (Advanced Analytical Technologies Inc. Fragment Analyzer) showed DNA fragments of variable length to be present in the genomic libraries. The curve shown presented here is for samples that had unquantifiable DNA using BR Qubit, suggesting that DNA is still present at very low concentrations. The two peaks (LM and UM) indicate lower and upper size standards, and the number at the top of the graph shows the average length (bp) of DNA strands in the genomic library. The panel on the right shows a smear analysis of the total DNA library in grey, and fragments size standards in blue.

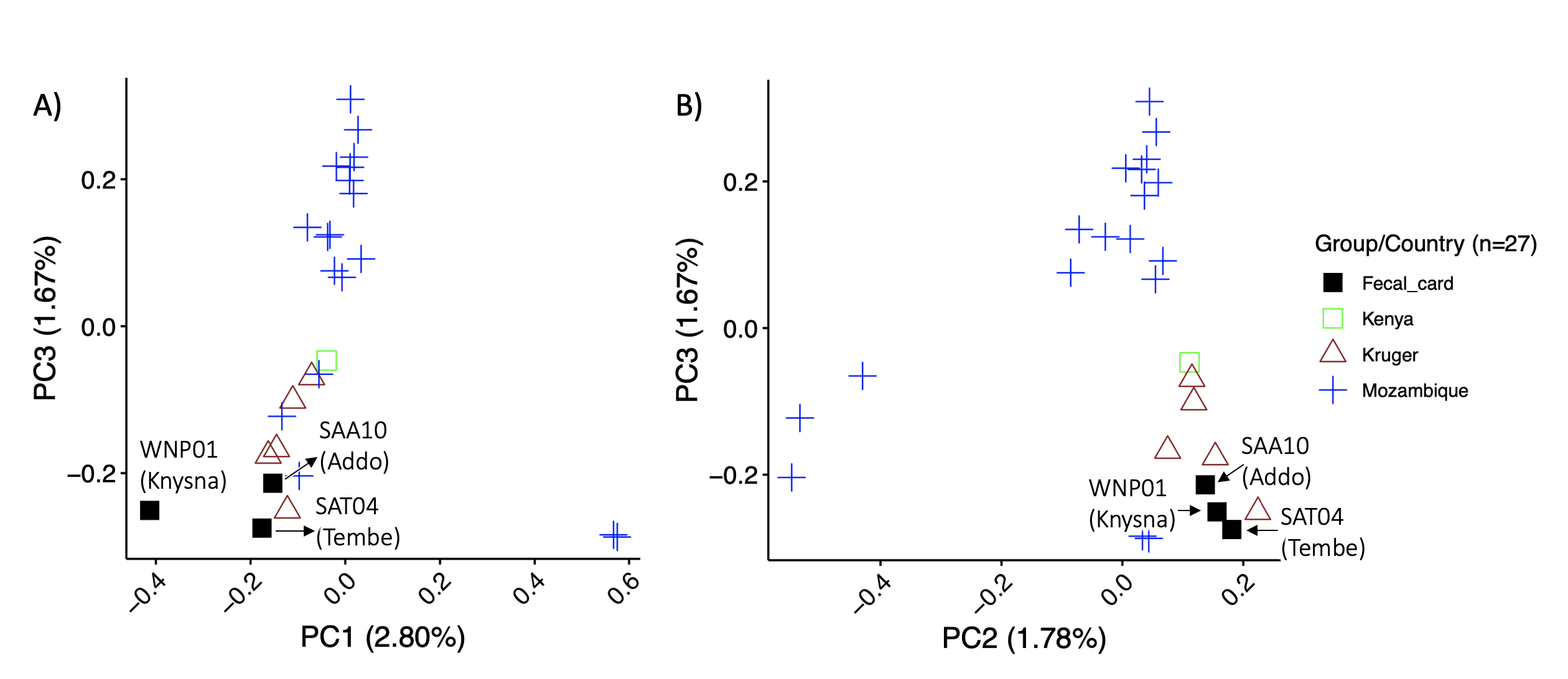

Supplementary Figure 7. Principal component analysis of nuclear genome-wide SNPs (single nucleotide polymorphisms) supported phylogeographic partitioning of elephants sampled via fecal cards and previously sequenced georeferenced elephants. Elephants from South Africa (high quality DNA samples from Kruger and fecal card samples) clustered together in the PCA. Panel A shows the clustering pattern for PC1 and PC3, where elephant samples collected on fecal cards from Addo Elephant Reserve and Tembe Elephant Reserve group with geo-referenced individuals from Kruger National Park, while the Knysna elephant (WNP01) does not group with any other individual. Panel B shows the clustering pattern for PC2 and PC3, where all elephants sampled via fecal cards group together and with previously sequenced georeferenced elephants from Kruger National Park. Symbols and colors represent different geographic localities (legend to the right of the panel).

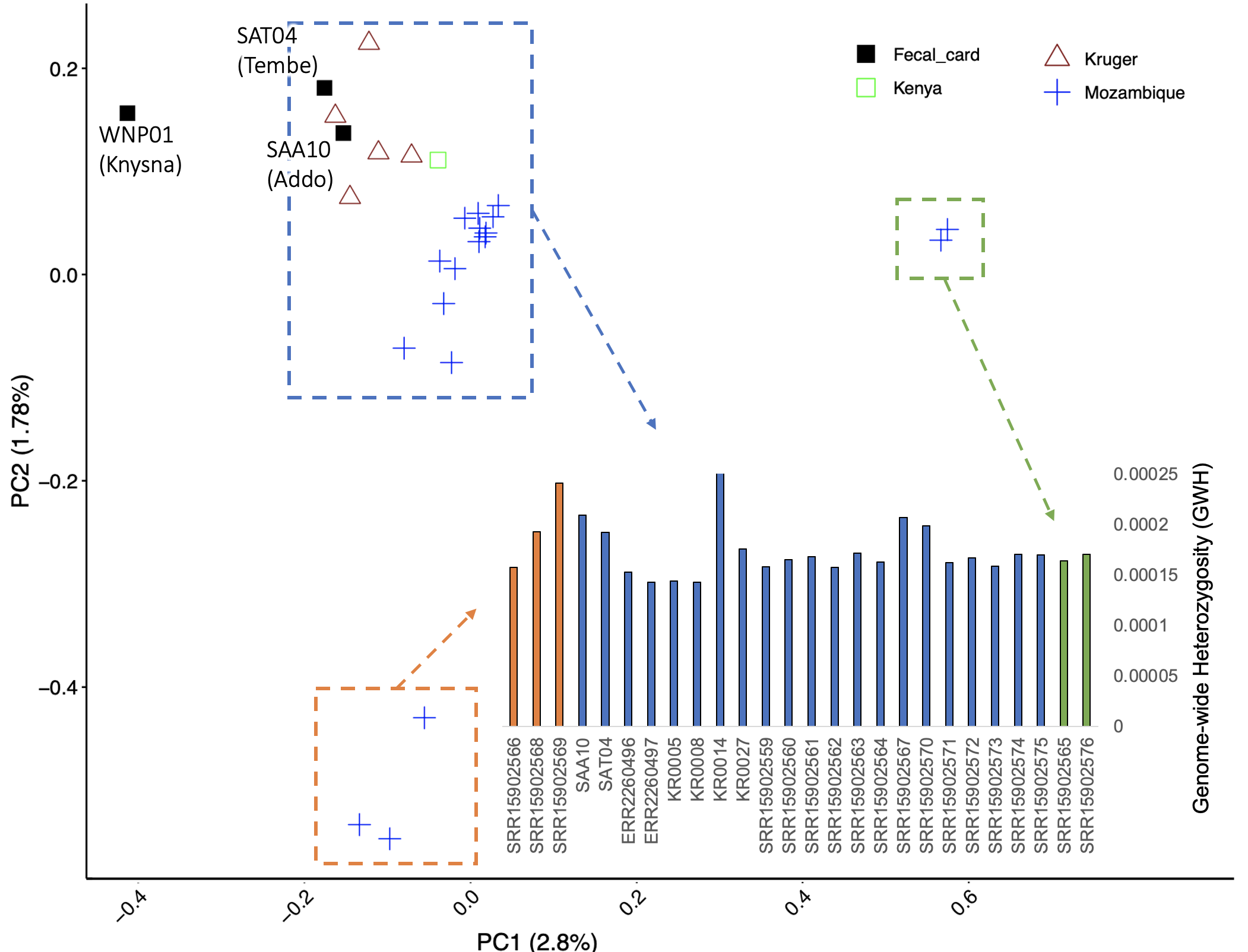

Supplementary Figure 8. Principal component analysis of nuclear genome-wide SNPs (single nucleotide polymorphisms; main graph) showed that elephants were not clustered based on genome-wide heterozygosity (GWH; inserted bar-graph in the lower-right of panel). Colored symbols represent different geographic localities (legend on the top-right of the panel), and each cluster of individuals are indicated by a hatched-line rectangle that has been colored to correspond with the GWH of individuals (indicated with arrows). The plot showing GWH includes the sample name or accession number of each elephant (Supplementary Table 1 contains additional information for each elephant).

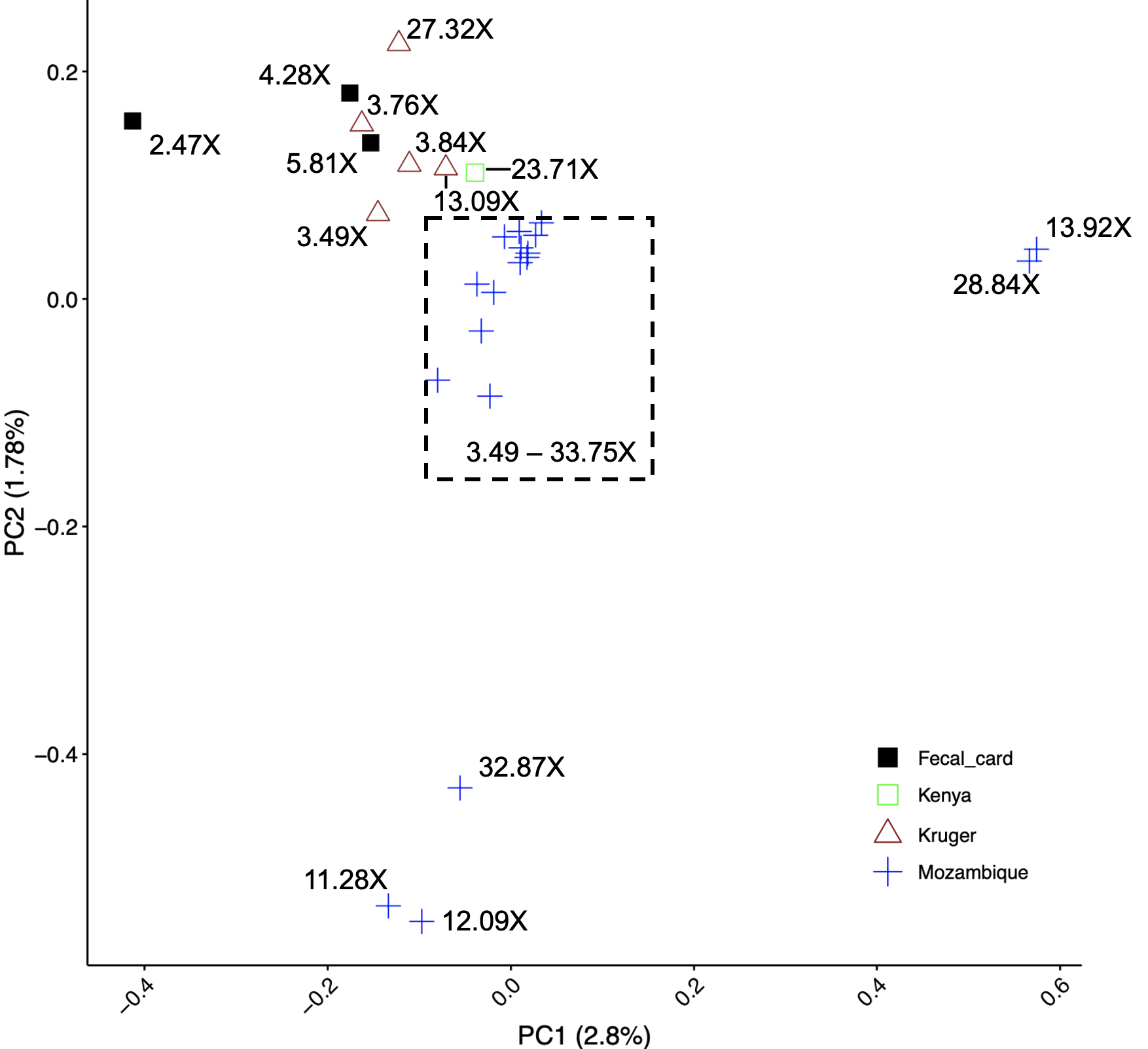

Supplementary Figure 9. Principal component analysis of nuclear genome-wide SNPs (single nucleotide polymorphisms) showed that elephants were not clustered based on genome sequence coverage (X-fold coverage of nuclear genome is indicated with text next to each symbol, or indicated as a range for a cluster of individuals within a hatched rectangle). Colored symbols represent different geographic localities (legend on the bottom-right of the panel).
